# A cross-cohort analysis of autosomal DNA methylation sex differences in the term placenta

**DOI:** 10.1101/2021.03.08.434471

**Authors:** Amy M. Inkster, Victor Yuan, Chaini Konwar, Allison M. Matthews, Carolyn J. Brown, Wendy P. Robinson

**Affiliations:** BC Children’s Hospital Research Institute, 950 W 28th Ave, Vancouver, Canada, V6H 3N1; Department of Medical Genetics, University of British Columbia, 4500 Oak St, Vancouver, Canada, V6H 3N1; Centre for Molecular Medicine and Therapeutics, 950 W 28th Ave, Vancouver, Canada, V6H 3N1; Department of Pathology & Laboratory Medicine, University of British Columbia, 2211 Wesbrook Mall, Vancouver, Canada, V6T 1Z7

**Keywords:** DNA methylation, placenta, sex as a biological variable, sex differences, microarray, Illumina 450K array, epigenetics, pregnancy

## Abstract

**Background:** Human placental DNA methylation (DNAme) data is a valuable resource for studying sex differences during gestation, as DNAme profiles after delivery reflect the cumulative effects of gene expression patterns and exposures across gestation. Here, we present an analysis of sex differences in autosomal patterns of DNAme in the uncomplicated term placenta (n=343) using the Illumina 450K array.

**Results:** Using a false discovery rate < 0.05 and a mean sex difference in DNAme beta value of > 0.10, we identified 162 autosomal CpG sites that were differentially methylated by sex, and that replicated in an independent cohort of samples (n=293). Several of these differentially methylated CpG sites were part of larger correlated regions of differential DNAme, and many also exhibited sex-specific DNAme variability. Although global DNAme levels did not differ by sex, the majority of significantly differentially methylated CpGs were more highly methylated in male placentae, the opposite of what is seen in differential methylation analyses of somatic tissues. Interestingly, patterns of autosomal DNAme at these significantly differentially methylated CpGs organized placental samples along a continuum, rather than into discrete male and female clusters, and sample position along the continuum was significantly associated with maternal age and newborn birthweight standard deviation.

**Conclusions:** Our results provide a comprehensive analysis of sex differences in autosomal DNAme in the term human placenta. We report a list of high-confidence autosomal sex-associated differentially methylated CpGs, and identify several key features of these loci that suggest their relevance to sex differences observed in normative and complicated pregnancies.

## BACKGROUND

Sex is a key variable that influences biological systems from the level of the cell to the level of the organism. Considering cells, tissues, and organs, biological sex is typically defined by sex chromosome complement, which largely corresponds with the gonadal sex of the organism (1). Biological sex is of particular importance in the study of human pregnancy and prenatal development as male fetal sex is a risk factor for several pregnancy complications including preterm birth, intrauterine growth restriction, and maternal gestational diabetes (2–6). Sex differences during prenatal development are likely affected by sex differences in the placenta, the organ critical for regulating growth and development of the embryo/fetus throughout gestation. Except in rare cases, placental cells harbor the same sex chromosome complement as the fetus, and sex differences in placental function, for example placental response to infection and stress, could contribute to sex differences in fetal growth and development (5,7,8).

Placental DNA methylation (DNAme) data provide valuable resource for studying sex differences during gestation, as DNAme profiles after delivery reflect the cumulative effects of gene expression patterns and exposures across gestation. In any tissue, when evaluating sex-specific DNAme both autosomal and X chromosomal loci should be considered. Sex differences in X chromosome DNAme are extensive and expected, as DNAme plays a key role in the process of X chromosome inactivation (XCI), by which the one X chromosome in female cells becomes epigenetically silenced via the accumulation of heterochromatic marks (9, 10). In contrast, the extent to which sex differences in autosomal DNAme patterns exist is less clear. Initial reports of sex-specific autosomal DNAme based on microarray data were later deemed false positives, attributed to probes determined *in silico* to have high sequence affinity to bind multiple genomic regions, many of which map to both autosomal and X or Y-linked loci (11, 12). The removal of CpG sites measured by such cross-hybridizing probes is now commonplace prior to most analyses of DNAme data, but rarely are sex differences at the remaining autosomal CpGs investigated. As a result, literature investigating sex differences in placental autosomal DNAme and gene expression patterns is sparse. However, the handful of studies conducted using placentae from uncomplicated pregnancies suggest that the placenta harbors an appreciable number of autosomal loci with sex-specific DNAme profiles (13, 14) and that a large proportion (potentially up to 60%) of sex-differentially expressed placental genes are autosomal (15, 16).

In the context of pregnancy, autosomal epigenome-wide association studies are routinely conducted to investigate the effects of factors such as disease phenotypes including preeclampsia and intrauterine growth restriction, and environmental exposures such as pollution or maternal smoking, on the placental epigenome (17–22). Understanding how biological sex is associated with autosomal DNAme is a relatively unexplored facet of prenatal epigenetic research, and may shed light on the factors contributing to sex differences observed in growth and development throughout gestation. This study seeks to comprehensively characterize sex differences in the uncomplicated, full-term (> 37 weeks of gestation) placental epigenome, with the aim of establishing a baseline of sex differences observed in the placental epigenome.

## METHODS

### Datasets

For the discovery cohort, placental Illumina Infinium HumanMethylation450 (450K) DNAme data obtained from liveborn deliveries were compiled from seven publicly available datasets from five North American cohorts (n=585). Datasets were selected on the basis of available infant birthweight and self-reported maternal ethnicity corresponding to one of three major ethnic identities (Black/African/African American, Asian/East Asian, European/White). The datasets compiled in this step include GSE73375 (n=9, North Carolina, USA) (23), GSE75428 (n=289, Rhode Island Child Health Study, Rhode Island, USA) (24), GSE98224 (n=9, Toronto, Canada) (25), GSE74738, GSE100197, GSE108567, and GSE128827 (n=34, all Epigenetics in Pregnancy Complications Cohort, Vancouver, Canada) (26–29). These data were utilized as described in Yuan et al. 2019, to generate PlaNET, the Placental DNAme Elastic Net Ethnicity Tool, which estimates metrics of genetic ancestry from placental DNAme datasets (28).

For replication analyses, an independent North American dataset was used, the New Hampshire Birth Cohort Study, New Hampshire, USA (n=293), GSE71678. Samples from the replication dataset were kept independent from the discovery cohort during preprocessing and analysis, analogous methods were used in both cohorts.

### Verification of sample sex and identity

In both the discovery and replication cohorts, two approaches were used to verify that the sex chromosomal complement of each sample corresponded to the sample sex as annotated in the metadata. First, samples were subjected to hierarchical clustering on DNAme β values (a metric ranging from 0 to 1 reflecting percent methylation) from only CpGs mapping to the X and Y chromosomes (n=11,648). Two major clusters corresponding to male (XY) and female (XX) samples were observed. Subsequently, samples were clustered using only the β values associated with five CpGs in the X inactivation centre, which reflects the X chromosome complement as these probes are proportionally methylated to the number of chromosomes silenced by XCI (10). Again, two major groups of samples were observed: samples annotated as female fell into one cluster that we deemed “XX”, while samples annotated as male fell into the second, deemed the “X” cluster as this check gives no information about Y chromosome complement. Samples were confirmed to be male or female if both sex chromosome clustering checks agreed with sex as annotated in the sample metadata.

Additionally, all samples were confirmed to be genetically unique using the “ewastools” R package (30). Two samples were found to be genetic duplicates in the replication cohort; after confirming by clustering on beta values from the 65 rs probes, these samples were both excluded from downstream analyses. Following sex and identity verification, the 65 rs genotyping probes on the 450K array and 11,648 CpGs mapping to the X and Y chromosomes were removed from both the discovery and replication datasets.

### Data processing and ancestry estimation

All samples from the discovery cohort (n=585) were subjected to routine filtering and normalization as described in Yuan et al. 2019 (28). Briefly, CpGs removed included those targeted by non-specific probes (31, 32), known placental-specific non-variable CpGs (range of β values < 0.05 between the 10^th^-90^th^ centile in all samples in this cohort) (33), and those with poor quality data (detection P value > 0.01 or bead count < 3 in more than 1% of samples) (34). Data were normalized by the normal exponential out-of-band (noob) and beta mixture quantile (BMIQ) normalization methods from the R packages “wateRmelon” and “minfi”, respectively (35). Samples were assigned values in three continuous ancestry coordinates, reflecting the probability of each sample being similar to 1000 Genomes Project populations of African, Asian, and European descent (28). Following the development of PlaNET, probes targeting polymorphic loci were also removed from this cohort (31, 32), as were samples born before term (<37 weeks gestation) and/or affected by preeclampsia; this left 324,104 autosomal CpGs in 341 samples available for sex-specific DNAme analysis.

The replication cohort was processed similarly; first, samples were restricted to those born >37 weeks of gestation, no other pregnancy complications affected these samples according to the metadata. Next, ancestry was estimated using PlaNET, all samples were found to be predominantly of European ancestry, as reported in the original publication (21), and filtering was conducted identically to the discovery cohort. To correspond with the original publication of this dataset, functional and noob normalization were performed (21). After filtering and normalization, 341,939 autosomal CpGs in 293 samples remained for replication analyses.

### Global sex-specific DNAme profile analyses

In order to study sex differences in global DNAme profiles, mean DNAme β values by sex at 324,104 filtered autosomal loci and 12,329 additional CpGs annotated to autosomal Alu and LINE1 repetitive elements were investigated by non-parametric Kruskal-Wallis tests. CpGs in repetitive regions were identified by the overlap of Illumina probe locations and the UCSC hg19 RepeatMasker track (36). The non-filtered dataset (n=473,929 CpGs, 341 samples) was used for this analysis as exclusion of CpGs in repetitive elements is a standard preprocessing step in EWAS. Additionally, sex differences in the proportions of fully methylated (β > 0.99), highly methylated (β > 0.90), lowly-methylated (β < 0.10), and unmethylated (β < 0.01) autosomal CpG sites in the filtered dataset were evaluated by Wilcoxon rank-sum tests.

To test whether placental cell composition differed by sex, the relative proportions of trophoblast, syncytiotrophoblast, stromal, endothelial, Hofbauer, and nucleated red blood cells in all discovery cohort samples were estimated using the reference-based method implemented in PlaNET (28). Sex differences in cell type proportions were evaluated using a linear model with cell type proportion as the outcome variable, adjusting for gestational age, dataset location of origin (location), and PlaNET-inferred ancestry coordinates 2 and 3, included as continuous additive covariates in the linear model. As the predicted ancestry coordinates are compositional, coordinates 2 and 3 were selected as they had the highest mean values across samples of the 3 coordinates.

### Identification of site-specific sex-associated autosomal DNAme

Sex-specific autosomal differentially methylated positions (DMPs) were identified in the discovery cohort by linear modelling on log-transformed β values (M-values) using the limma package in R (37); gestational age, dataset of origin (location), and PlaNET coordinates 2 and 3 were included as covariates. Sex differences in DNAme β values at each individual CpG site were calculated as Δβ = Average Male β – Average Female β. Multiple test correction was performed with the Benjamini-Hochberg false-discovery rate (FDR) method. In the replication cohort, a similar model was used except that PlaNET-inferred ancestry coordinates were not included in the linear model, as the PlaNET-estimated ancestry of all 293 samples was extremely homogeneous (predominantly European), see Supplementary Figure 1. For all downstream analysis, DMPs were considered replicated if they satisfied the following criteria in GSE71768: FDR < 0.05 and Δβ > 0.05 in the same direction as the discovery cohort.

### BLAST analysis for cryptic sex chromosome-associated DNAme

Next, DMPs were evaluated for evidence of probe cross hybridization to other genomic loci, especially the sex chromosomes. As described in Chen et al. 2011, command-line nucleotide BLAST (blastn) was performed on the 50 nucleotide probe sequences for all replicated DMPs, searching against four versions of hg19 (*in silico* bisulfite converted fully methylated and fully unmethylated, both forward and reverse complement) (12). BLAST results representing the intended hybridization targets per the Illumina 450K array manifest were removed from the list of results, remaining sequences were considered non-specific with a BLAST match of at least 40 sequential nucleotides and a nucleotide match at position 50. Non-specific DMPs with matches on the X or Y chromosome were removed from the list of replicated DMPs used in downstream analysis.

### Gene ontology analyses

An enrichment analysis of biological process terms from the Gene Ontology collection was conducted on the genes associated with DMPs that replicated in our second cohort using the “gometh” and “goregion” functions from missMethyl, which accounts for potential bias from Illumina arrays measuring DNAme at multiple CpGs per gene (38). Genes associated with all 324,104 autosomal CpGs tested for differential DNAme by sex were used as the background set. Gene ontology terms satisfying a threshold of FDR < 0.05 were considered significantly enriched in the geneset associated with the top DMPs.

### Proximity to transcription factor binding motifs

DMPs were examined for enrichment in proximity (100bp window with the CpG of interest at the centre) to transcription factor binding motifs from Homo sapiens Comprehensive Model Collection (HOCOMOCO) version 11 as compared to the background list of 324,104 filtered autosomal CpGs; this analysis was conducted using the CentriMo tool for local enrichment analysis from the Multiple Em for Motif Elicitation (MEME) Suite browser tool (39–41). In addition, enrichment for androgen receptor (AR) and estrogen receptor (ER) α and β binding sites within a 100bp window with the CpG of interest at the centre was tested as compared to the input filtered autosomal probe set. Genomic coordinates of AR and ER binding sites were obtained from Wilson et al. 2016 and Grober et al. 2011 (42, 43); enrichment was assessed using exact goodness-of-fit tests.

### Relationship between sex-specific DNAme and differentially expressed genes by sex

Preprocessed and normalized placental gene expression data was downloaded for GSE75010, collected with the Affymetrix Human Gene 1.0 ST Array (44). Non-preeclamptic samples from this cohort born at or after 37 weeks of gestation were selected for our analyses (n=34, 47% female). The 65 genes covered by the Affymetrix array that overlapped replicated DMPs were tested for differential expression by sex via linear modelling, adjusting for maternal hypertension (yes/no), self-reported ethnicity, and gestational age at birth. Genes were considered differentially expressed by sex at nominal significance (p < 0.10).

### Further characterization of differentially methylated CpG sites

Differentially methylated genomic regions (DMRs) were identified using the R package DMRcate with lamba=1000 and C=2 considering all 324,104 autosomal CpGs (45). DMRs were considered significant at an FDR < 0.05 if comprised of at least 3 CpG sites with a mean Δβ across the region of > 0.05 in either direction, calculated as Δβ = Average Male β – Average Female β. A lower Δβ was tolerated in this analysis as it was a regional average.

DNAme loci that were differentially variable methylated positions by sex (DVPs) were identified using the iEVORA function from the “matrixTests” R package, with all 324,104 autosomal CpGs as input (46). This method was selected as it ranks selected features by differential mean DNAme t-test p values to decrease the likelihood of identifying differentially variable positions driven by single sample outliers (46).The cut and cutBfdr thresholds used were 0.05 and 0.001, respectively.

### Sex continuum analysis

For a subset of samples from the Vancouver cohort (datasets GSE74738, GSE100197, GSE109567, GSE12887, n=34, 53% female), we had access to extended clinical information beyond the demographics presented in Table 1. These samples were selected from the discovery cohort to investigate associations between various clinical phenotypes and sample score along the first principal component of the 162 DMPs. Relationships between demographic variables and sample scores along the first principal component (PC1) were assessed by linear modelling, with PC1 score as the outcome variable and each clinical variable used as a predictor in independent models. Categorical variables informative across all samples included 450K array row, chip, and batch, positive maternal serum screen, and delivery type. Continuous variables informative across all samples included gestational age, maternal body mass index, maternal age, birth weight, birthweight standard deviation z-score corrected for infant sex and gestational age, processing time between delivery and placental sampling, and the estimated proportion of major placental cell types estimated using the PlaNET algorithm.

**Table 1.** Demographic characteristics of discovery and replication cohorts.

|  | Discovery |  |  | Replication |  |  |
| --- | --- | --- | --- | --- | --- | --- |
|  | Female (n=177) | Male (n=164) | p value | Female (n=137) | Male (n=156) | p value |
| **Gestational Age** |  |  |  |  |  |  |
| Weeks | 39.0 (± 1.1) | 39.1 (± 0.9) | 0.53 | 39.6 (± 1.1) | 39.7 (± 1.0) | 0.28 |
| **Condition** |  |  |  |  |  |  |
| Healthy term | 100 | 100 | 0.02 | 114 | 125 | 0.10 |
| SGA | 37 | 45 |  | 9 | 4 |  |
| LGA | 40 | 19 |  | 13 | 24 |  |
| **PlaNET<sup>†</sup> Scores** |  |  |  |  |  |  |
| Coordinate 1 (mean (SD)) | 0.10 (± 0.25) | 0.07 (± 0.23) | 0.27 | 0.0009 (± 0.0016) | 0.0012 (± 0.0018) | 0.000024 |
| Coordinate 2 (mean (SD)) | 0.11 (± 0.26) | 0.05 (± 0.14) | 0.37 | 0.0038 (± 0.0270) | 0.0032 (± 0.0103) | 0.011 |
| Coordinate 3 (mean (SD)) | 0.78 (± 0.36) | 0.88 (± 0.27) | 0.23 | 0.9951 (± 0.0279) | 0.9956 (± 0.0111) | 0.001 |

*SD refers to standard deviation, SGA and LGA refer to small (<10<sup>th</sup> centile) and large (>90<sup>th</sup> centile) birthweight for gestational age within each sex, as assigned by the original publications. \*p values are from Wilcoxon rank-sum tests for continuous and Fisher's exact test for categorical variables. <sup>†</sup> PlaNET coordinate 1 is associated with African ancestry, coordinate 2 with Asian ancestry, and coordinate 3 with European ancestry (28).*

## RESULTS

### Genome-wide measures of DNAme do not differ by placental sex

To investigate whether female (XX) and male (XY) term placentae had different global DNAme profiles, we tested for an association between sex and genome-wide DNAme using all autosomal CpGs in the filtered dataset (n=324,104). Neither the overall mean β value nor the proportions of highly methylated (> 0.90), fully methylated (> 0.99), or lowly methylated (< 0.10) and unmethylated (< 0.01) autosomal CpGs differed by sex in this cohort. Repetitive elements are frequently interrogated as surrogates for global DNAme as they comprise roughly 30% of all genomic nucleotides, as well as 30% of CpG dinucleotides (47). However, sex was not significantly associated with mean DNAme at Alu or LINE1 repetitive elements (Kruskal Wallis p > 0.05).

DNAme profiles obtained from bulk tissue such as whole blood, buccal swab, and placenta reflect the proportion-weighted composite methylomes of all contributing cell types. When studying bulk tissue DNAme, it is important to consider how sampling procedures and/or biology may alter relative cell type proportions in a biological sample, and how this may be reflected in results (48). We used the reference-based PlaNET algorithm to estimate the relative proportions of major placental cell types (trophoblasts, syncytiotrophoblasts, stromal cells, Hofbauer cells (placental macrophages), and endothelial cells), and found that the relative cell type proportions did not differ by placental sex (Figure 1).

**Figure 1.**
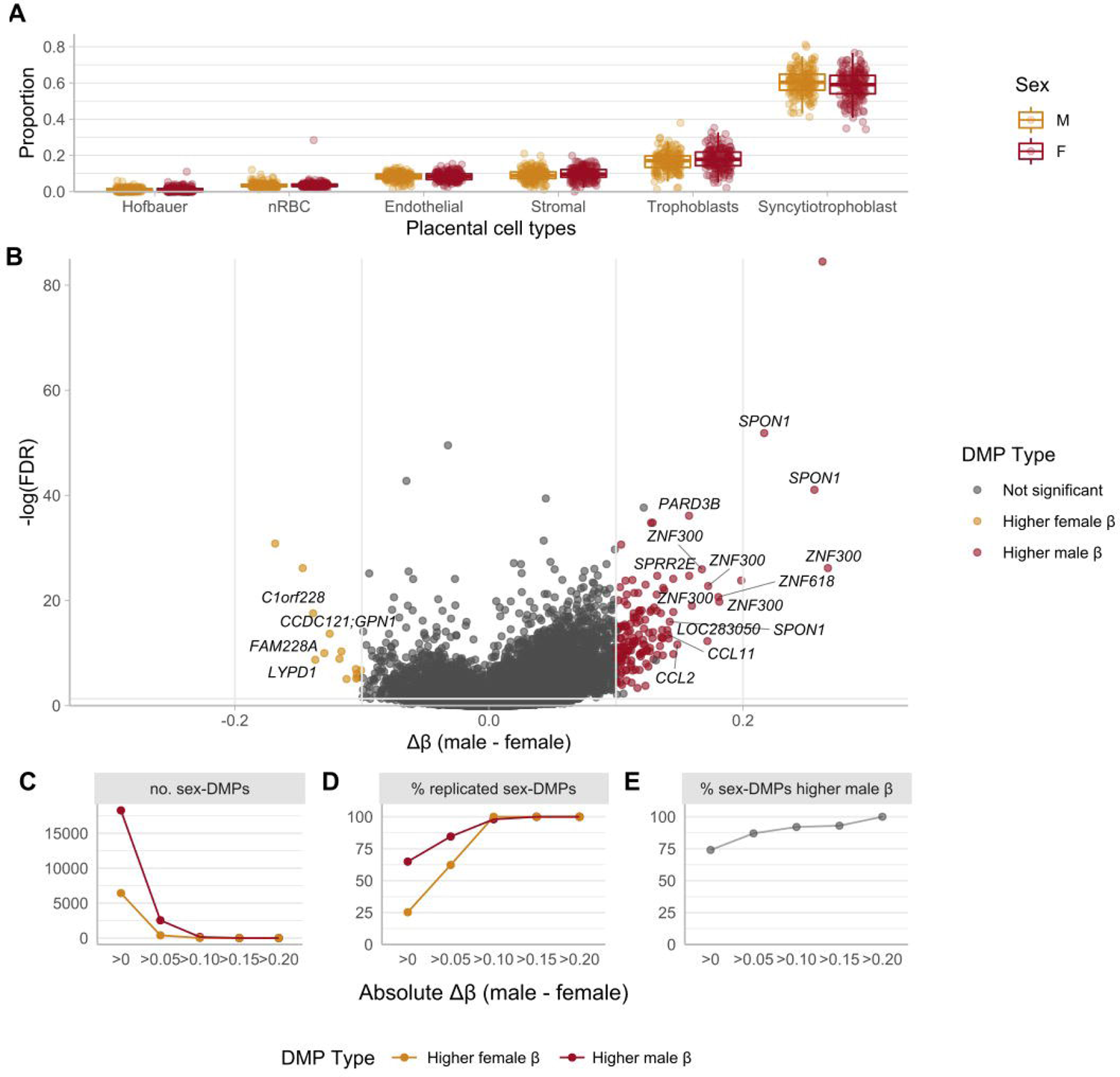
Sex differences in autosomal DNAme patterns by placental sex. **(A)** Estimated cell type proportions by sex in the discovery cohort, estimated using the R package PlaNET. Cell type proportions do not significantly differ by sex (p>0.05). **(B)** Volcano plot of all 324,104 autosomal CpG sites in the discovery cohort. Thresholds of statistical and biological significance are depicted by horizontal (FDR < 0.01) and vertical (Δβ > 0.10) intercepts. Significantly differentially methylated autosomal CpG sites by sex (FDR < 0.01, Δβ > 0.10) are highlighted in colour to indicate direction of sex-biased DNAme. CpG sites in yellow have significantly higher average male DNAme at these thresholds, red sites exhibit higher female DNAme. CpG sites not significantly differentially methylated by sex at these thresholds are in grey. Each point represents a single CpG site, Δβ= β_avgmale_-β_avgfemale_. The most differentially methylated CpG sites are annotated with associated genes names. **(C)** The number of differentially methylated (FDR<0.05) CpG sites at various Δβ thresholds; DMPs that are more highly methylated in male samples are indicated in red, DMPs more highly methylated in female samples are indicated in orange. **(D)** Percentage of DMPs at various Δβ thresholds that replicate (FDR<0.05, Δβ same direction) in GSE71678, colored by sex with higher DNAme. **(E)** For all DMPs at the Δβ thresholds considered, the percentage of DMPs with higher male DNAme.

### Male placentae show higher DNAme at a subset of autosomal loci

A linear model was fitted on M-values to test for differential DNAme by sex in the filtered dataset (n=324,104 autosomal CpGs), adjusting for gestational age at birth, dataset, and inferred genetic ancestry. The number of sex-associated CpG sites at various statistical (FDR) and biological (Δβ) thresholds were considered to evaluate the extent to which autosomal DNAme profiles in the placenta are affected by sex (Table 2).

**Table 2.** Results of linear modelling for sex-specific autosomal DNAme.

| | $\Delta\beta > 0$ | $\Delta\beta > 0.05$ | $\Delta\beta > 0.10$ | $\Delta\beta > 0.20$ |
| --- | --- | --- | --- | --- |
| **FDR < 0.05** | 24,715 (0.74) | 2,942 (0.87) | 166 (0.92) | 4 (1.00) |
| **FDR < 0.01** | 14,108 (0.80) | 2,682 (0.88) | 166 (0.92) | 4 (1.00) |

*Number of significantly differentially methylated autosomal positions at various statistical and biological thresholds are shown. FDR indicates the Benjamini-Hochberg false discovery rate, $\Delta\beta$ refers to the difference in DNAm $\beta$ value (male-female) and indicates the biological effect size between the sexes. The numbers in brackets indicate the proportion of sites at each threshold level that are more highly methylated in male placentae.*

To focus on CpGs more likely to be reproducible in future studies (27), CpGs were considered significantly differentially methylated by sex if they satisfied an FDR < 0.05, and an absolute Δβ > 0.10 between males and females. In total, 166 sex-associated differentially methylated positions (DMPs) fit these stringent criteria, of which 92% were more highly methylated in male samples than in females, a pattern that was observed across all thresholds considered (Figure 1, Table 2). See Supplementary Table 1 for the results of all investigated autosomal CpGs.

We hypothesized that some DMPs may contribute to larger regions of correlated sex-specific DNAme, as several of the DMPs overlapped the same genes and genomic regions. To test this, we performed differentially methylated region (DMR) analysis in the discovery cohort.

Significant DMRs were identified based on criteria of FDR < 0.05, spanning at least 3 CpGs, and having a mean Δβ > 0.05 across the region, a lower Δβ was utilized in DMR analysis than in DMP analysis, as it reflected the average DNAme β of all composite CpGs. Eighty-seven DMRs comprised of 435 CpGs satisfied these criteria. The 87 DMRs ranged in size from 36 to 3306 base pairs (mean 890 base pairs) and were comprised of an average of 5 CpGs per DMR; these regions were on average 6.3% differentially methylated between the sexes. Of the 87 DMRs, 29 (33%) included one or more of the 166 identified DMPs, and conversely, 46 of the 166 DMPs (28%) were part of DMRs. It is likely that most of these DMPs are part of local regions of differential DNAme, but that the array coverage is not sufficient for their detection. Genes overlapping sex-specific DMRs included several from the chemokine ligand CCL family (2, 11, 13), the keratin KRT family (6, 74), the LCE family (1B, 6A), the SPRR family (1A, 2A, 2C, 4), and the ZNF family (423, 300), including *ZNF300* and *ZNF423*, see Figure 2. *SERPINA6* overlapped a DMR more highly methylated in male samples. For a list of all identified DMRs, see Supplementary Table 2.

**Figure 2.**
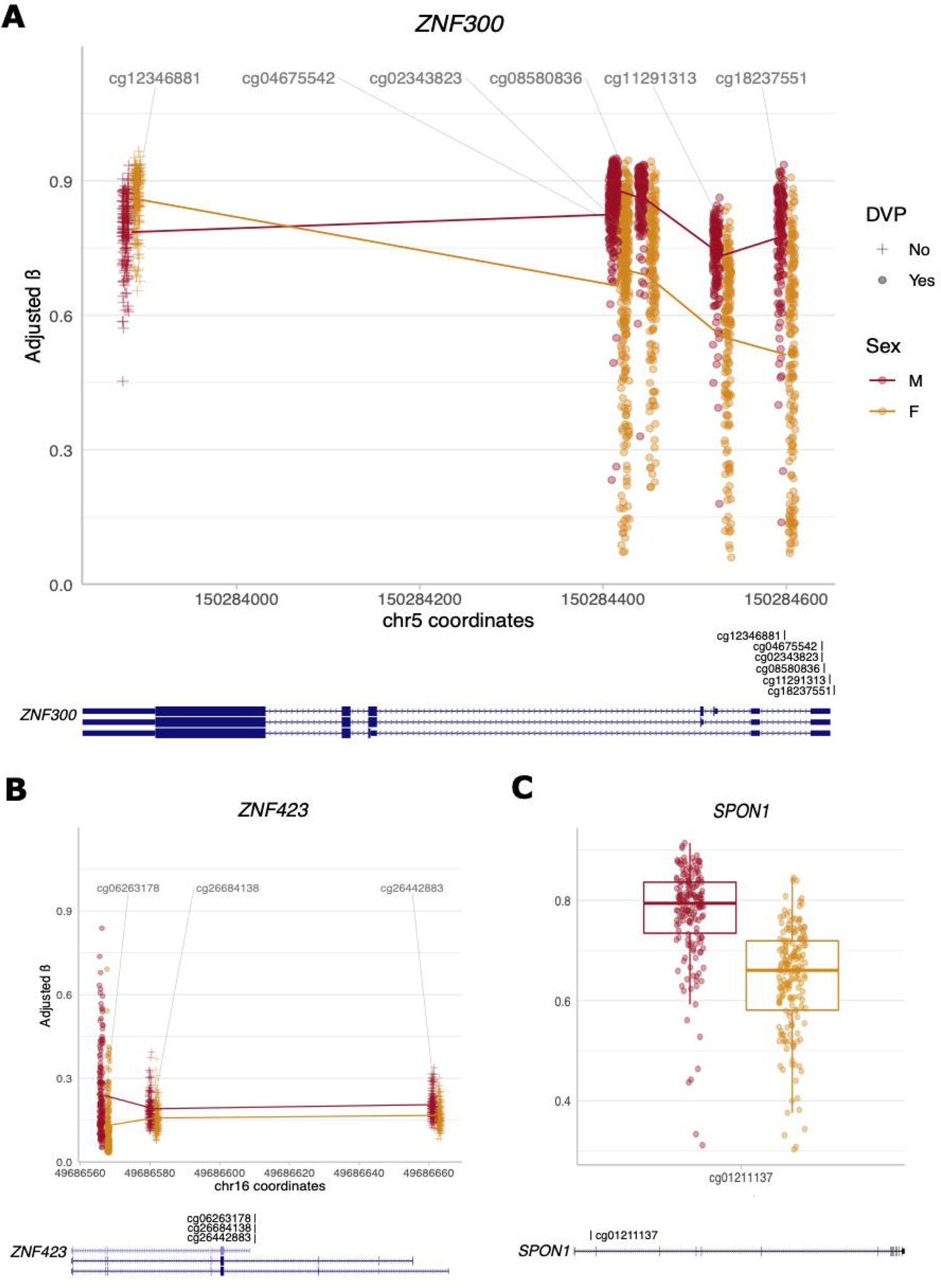
Scatterplots of sex-differentially methylated regions and probes in key genes. **(A)** Differentially methylated region spanning 5 CpGs in ZNF300 in chromosome 5, male samples are indicated in red, females in orange; the CpG coordinates along chromosome 5 are indicated on the X axis, while DNA methylation β values for each sample are plotted along the Y axis. CpG sites that are also significantly differentially variable (DVP) are indicated by circular scatter points. Below is the gene model from the UCSC Genome Browser track with the CpG positions indicated. **(B)** A differentially methylated region in ZNF423, coordinates along chromosome 16 are indicated on the X axis. **C)** A significantly differentially methylated CpG site in the gene body of SPON1, this site overlaps an estrogen receptor β binding site.

### More highly variable DNAme loci in female placentae

In addition to differences in mean DNAme at individual CpGs (DMPs), the DNAme variability may also vary by sex (DVPs). We undertook a DVP analysis in this study as part of a comprehensive characterization of sex differences in the placental DNA methylome, as to our knowledge no previous placental DVP studies have been reported, including related to sex. Differential variability in DNAme is an intriguing molecular feature often identified in cancer, which has many molecular correlates to successful placentation (49). A total of 3,148 significant (FDR < 0.001) DVPs were identified between the sexes, the majority of which were more highly variable in female samples than in male samples (n=3,170, 82%). Although no biological processes were significantly overrepresented in the gene set associated with these DVPs, the 6 nominally significant (p < 0.05) biological processes associated with DVPs between the sexes were related to cornification and keratinization, as well as neurological processes such as axon guidance, cerebral cell migration, glial cell-derived neurotrophic factor receptor signaling, negative regulation of neuron apoptosis, and dopamine uptake in synaptic transmission. Interpreting gene ontology enrichment analysis results in the placenta is difficult, though, as functional gene annotations provided in public databases are not placenta-specific, and genes in the placenta may function differently than in other tissues. Additionally, 19 of the DVPs were also DMPs (FDR < 0.05 and Δβ > 0.10) including CpG sites in the *SPON1* gene, see Figure 2. The results of the differential variability analysis are available in Supplementary Table 3.

### Replication of sex differences in DNAme

In EWAS studies, it is important to evaluate the robustness of any findings in an independent dataset to increase the likelihood of true positive findings. For replication, linear modelling was conducted to identify DMPs by sex in an independently processed Illumina 450K dataset, GSE71678 (n=293, 47% female). Because differences in DNAme (Δβ) are related to both biological and technical variables, and can vary for technical reasons alone by as much as 0.03-0.05, we used a less stringent Δβ threshold to define replication (27, 33). Of the 166 DMPs identified in the discovery cohort, 98% (n=163) replicated at an FDR < 0.05 and Δβ > 0 in the same direction as observed in the discovery cohort, see Figure 1.

### Genomic cross-reactivity of probes underlying sex-specific DNAme

DNAme at CpGs targeted by Illumina’s DNAme microarray is measured by 50 nucleotide probes that may cross-hybridize to off-target autosomal and sex chromosomal loci, and therefore have the potential to yield false positive results for sex-specific autosomal DNAme (12). To exclude the possibility that the sex-specific autosomal DNAme observed in this study was the result of sex chromosome cross-reactivity, we BLAST-ed the probe sequences associated with the replicated 163 DMPs against the hg19 human reference genome. We assessed all BLAST results matching greater than 40 nucleotides of the probe body with >90% sequence identity, and overlapping the 50^th^ nucleotide position (the CpG, hybridization at this nucleotide required for single base extension and fluorescence detection). Chen et al. and Price et al. used similar criteria define potential cross-hybridization (31, 32), although we chose to tolerate sequence matches with gaps in the interest of discovering even low-probability cross-reactivity to the sex chromosomes, as other studies have shown that 50-mer microarray probes may cross-hybridize to unintended regions with as little as 75-80% sequence identity if as few as 14 contiguous nucleotides match (50).

At these thresholds, only one probe showed evidence for possible cross reactivity to the sex chromosomes: cg02325951, which underlies a CpG site in the gene body of *FOXN3.* In the ProbeSeqA target sequence for this probe, 43 nucleotides match a region on the p arm of the X chromosome, approximately 1kb upstream of *HSD17B10* (chrX: 53467618-53467660). As we could not confidently determine whether the sex-specific DNAme observed at this CpG could be attributed to the intended genomic target (chr14: 89878619-89878668), we chose to exclude this CpG from downstream analyses of replicated hits (Supplementary Figure 2). This probe has previously been reported to be differentially methylated by sex in the placenta (13, 14).

### Characterization of autosomal sex-specific DMPs

The remaining 162 replicated and BLAST-ed DMPs were subsequently investigated for biological meaning. We observed no enrichment of specific genomic region locations (gene bodies, promoters, intragenic regions), on any particular autosomal chromosome, or for their position relative to CpG islands (located in CpG islands, shores, or shelves). Gene ontology analysis revealed significant enrichment for 10 biological process terms, which could be largely divided into two categories, the first related to chemokines/chemotaxis and immune function (chemotaxis; eosinophil, monocyte, and lymphocyte chemotaxis; chemokine-mediated signaling; cellular response to interleukin-1), and the second related to epithelial barrier function (peptide cross-linking, keratinocyte differentiation, keratinization, and cornification).

### Association with gene expression and transcription factor binding sites

We then tested whether the 65 genes associated with the 162 DMPs displayed sex-biased expression patterns. Of these 65 genes, only 8 were significantly differentially expressed between male and female placentae (p < 0.10), however this cohort of uncomplicated term placentae was small and therefore lacked statistical power (n=34 samples, 47% female). One such differentially expressed gene was *ZNF300*, which harbored a promoter DMP more highly methylated in males, and was more highly expressed in female placentae. ZNF300 has been previously reported to be more highly expressed in 46,XX human placentas (16).

Changes to DNAme at transcription factor binding motifs genome-wide can affect the efficiency of TF binding, either positively or negatively depending on the transcription factor, and may thus interact with gene expression patterns (51). Binding motifs for six transcription factors were significantly overrepresented within 100 nucleotide windows around the top DMPs (adjusted P value < 0.05 and CentriMo E-value < 1). These included motifs for AHR, ATF3, GMEB2, ZBT14, and two binding motifs for the KAISO protein (encoded by *ZBTB33*), see Table 3. *ZBTB33* is located on the X chromosome (Xq24), while the other transcription factors are encoded by autosomal genes. These transcription factors *AHR, ATF3, GMEB2, ZBTB33,* and *ZBTB14* were confirmed to be robustly expressed in the term placenta using dataset GSE75010, all five were more highly expressed than the median expression log2 counts-per-million of all placentally-expressed transcripts.

**Table 3.**
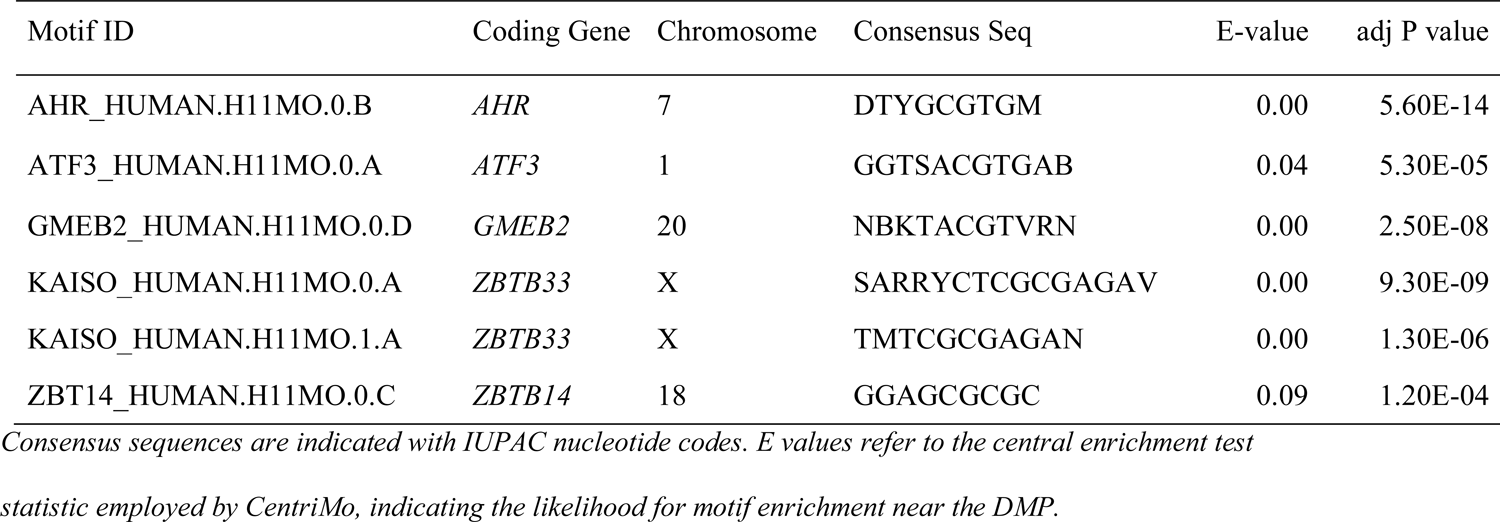
Transcription factor binding motifs overrepresented within 100bp of the top 162 DMPs.

We further tested whether the 162 DMPs were enriched for proximity to ER α and β and AR binding sites, as molecular sex differences can arise in general from the action of either sex chromosomes or sex hormones (1). We found no enrichment for ER α/β or AR binding sites within 200 base pair windows of the top DMPs, centered around the CpG of interest. However, there were two DMPs that overlapped AR and ER β binding sites, respectively. An intergenic CpG site on chromosome 8 overlapped an AR binding site, while a CpG site in the body of *SPON1* overlapped an ER β binding site, see Figure 2.

### Limited overlap of DMPs with previous studies

To contextualize the results of this study in the existing literature, we considered the overlap between DMPs identified as sex-associated in this study at an FDR < 0.05 (n=24,715) and two similar previous placental DNAme studies (Table 4) (13, 14). Due to different preprocessing criteria, and the fact that both previous studies relied on probes common to the 27k and 450k Illumina DNAme array platforms, not all identified DMPs in these studies were covered by probes in our dataset, and thus we restricted to comparing those that were. There was no overlap between DMPs found in our study and the 17 DMPs from Martin et al. (2017) (13), which only included preterm births <28 weeks of gestational age. However, 84 of the 335 (255) DMPs and 154/335 genes identified by Mayne et al. 2017 were also identified as part of our 166 DMPs (14).

**Table 4.**
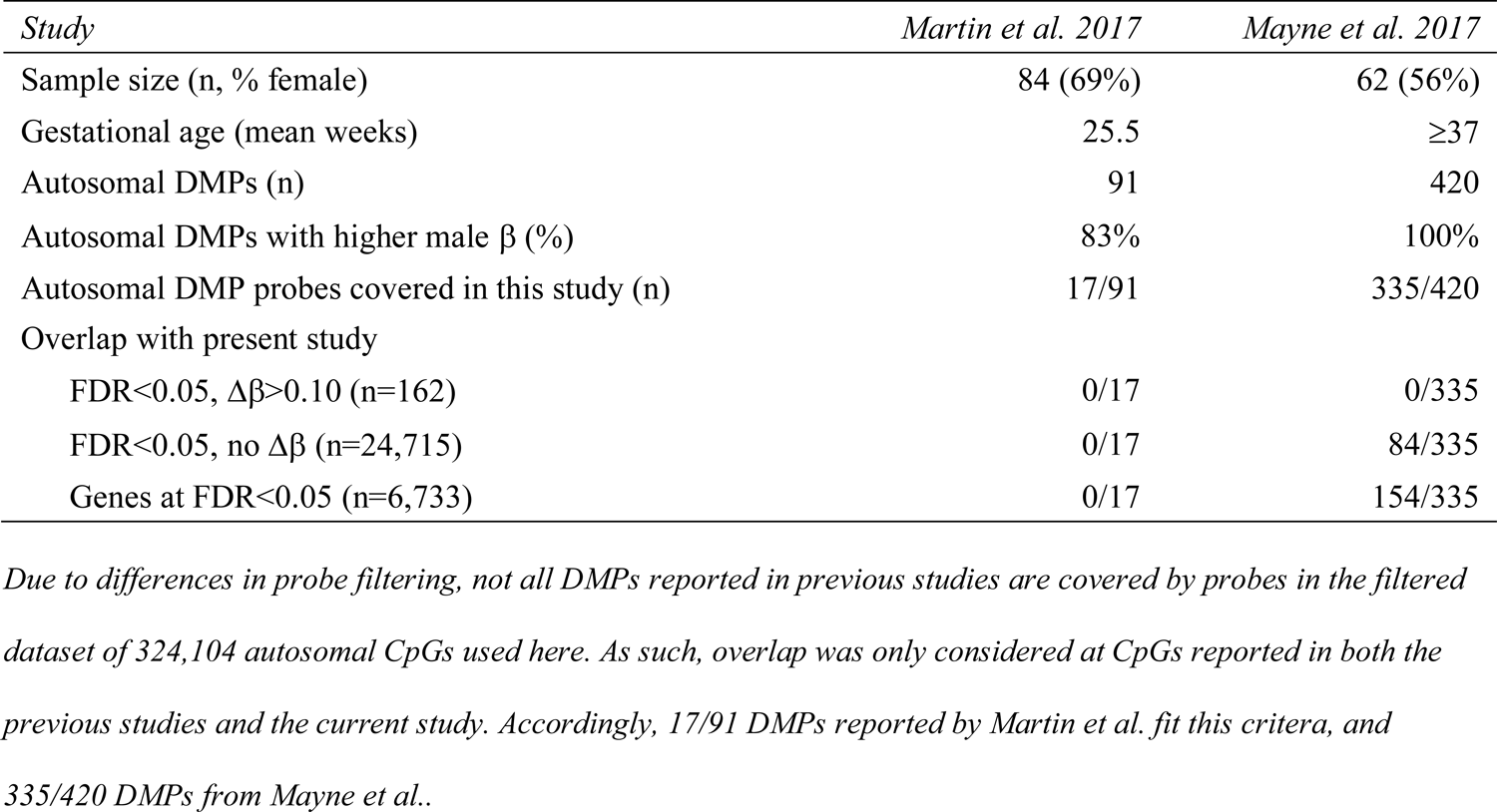
Overlap of placental autosomal differentially methylated CpGs reported in this study with previous literature.

### Combined effect of sex-specific DNAme at DMPs

To evaluate the combined effect of differential methylation at the 162 DMPs, we performed principal components analysis on the β values associated with these sites in all samples. Although both PC1 (37.1% variance) and PC2 (4.76% variance) were significantly associated with sample sex (ANOVA p < 0.05, respectively), rather than samples forming clearly delineated “male” and “female” clusters in PC space, they were instead distributed across PC1 in a continuum of sex, see Figure 3.

**Figure 3.**
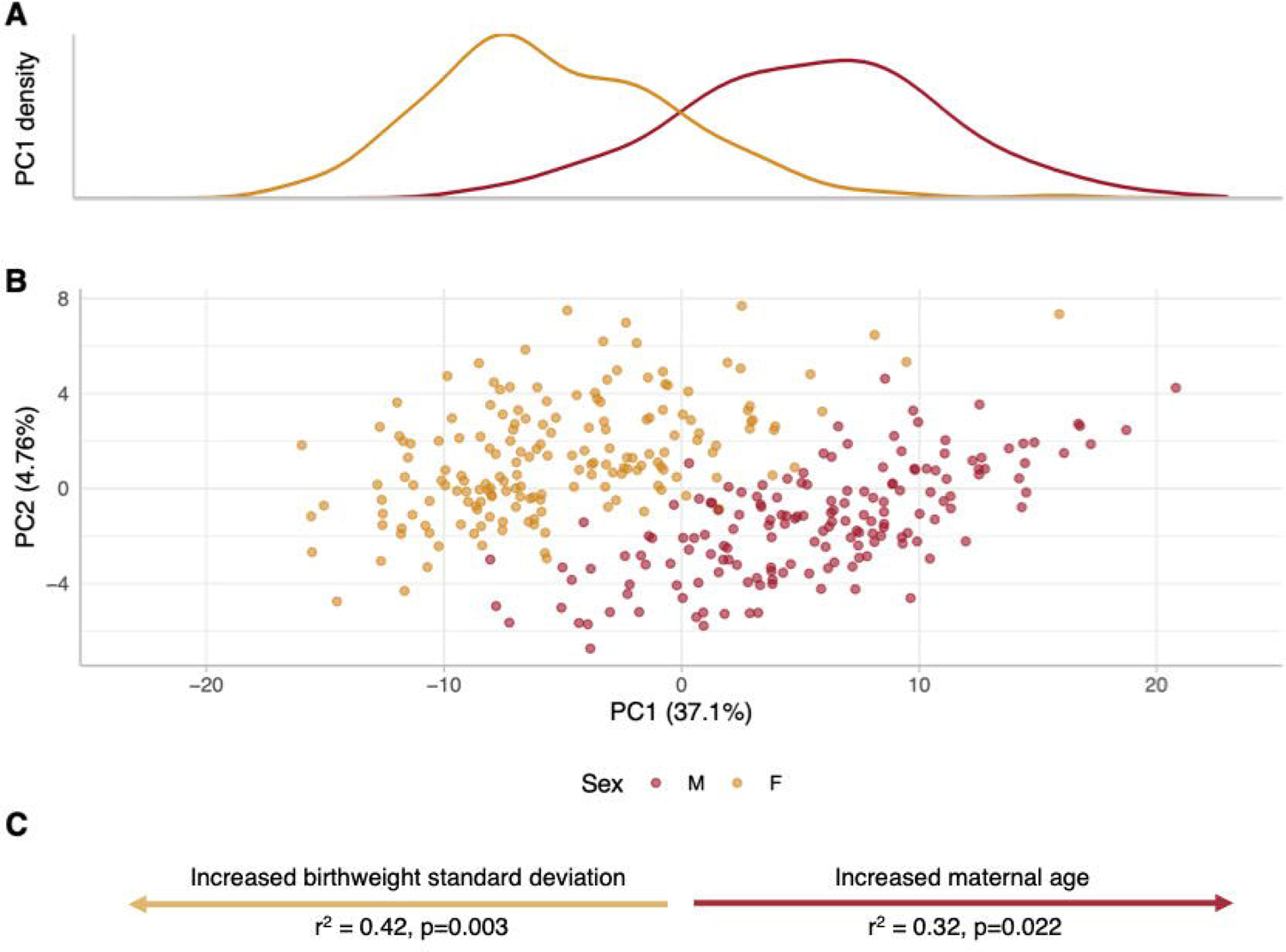
Principal components analysis of DNAme at the 162 significantly sex-differentially methylated CpGs. **(A)** Density plot of male (M, red curve) and female (F, yellow curve) samples along the first principal component. Principal components analysis was conducted on the discovery cohort based on the DNAme β values at the 162 sex-associated DMPs. **(B)** Scatterplot of principal component 1 versus principal component 2 of the discovery cohort., male (M) samples are plotted in red, females (F) in yellow. **(C)** Significant associations between clinical variables and the first principal component in a subset of 34 samples with extended metadata. R^2^ and p values are reported for each significant variable. Yellow arrow indicates female-only significant association, red arrow indicates male-only significant association. PC: principal component.

As sex biases are observed in the frequency and severity of many pregnancy complications, we hypothesized that a sample’s position along this continuum of sex (PC1) may be associated with sex-specific clinical features, such as birthweight. Using a subset of 34 samples from the discovery cohort for which we had extended metadata, we tested for a relationship between PC1 score and the following variables: positive maternal serum screen, delivery type, gestational age at birth, maternal body mass index, maternal age, birthweight standard deviation z-score adjusted for infant sex and gestational age, and the estimated proportion of major placental cell types estimated using the PlaNET algorithm. We also tested for associations with potential technical confounders including sample processing time after delivery and 450k array row, chip, and batch. When considering both sexes together, no clinical or technical characteristics informative across all samples were significantly associated with sample position along PC1. However, when stratifying analyses by sex, maternal age was significantly positively associated with PC1 score in males (higher maternal age in towards the male end of the continuum), while birthweight standard deviation was significantly positively associated with PC1 score in female samples (higher birthweight standard deviation toward the female end of the continuum).

We also leveraged the principal components analysis to further investigate the relationship between gestational age and sex chromosome complement with patterns of DNAme at the top differentially methylated loci. Twenty four second trimester and early third trimester samples (21-32 weeks), including three with 45,X chromosome complements were projected into the PCA space associated with the 162 differentially methylated CpG sites in the discovery cohort. The younger gestational age male and female samples still formed a sex continuum, but were localized to the top half of the plot, indicating that PC2 is associated with gestational age (p<2.2e-16). The 45,X samples were found to localize to the ‘male’ side of the first principal component within this cluster of younger GA samples, see Supplementary Figure 4.

## DISCUSSION

We undertook this study to identify the extent to which the human placenta exhibits true patterns of sex-specific autosomal DNA methylation, after rigorously accounting for probe cross-reactivity to the sex chromosomes. By compiling a dataset profiling DNAme in 341 term placentae, we identified, replicated, and analyzed the biological significance of sex-associated placental autosomal DNAme. There was no evidence for sex differences in placental cell type proportions underlying autosomal DNAme sex differences, nor was there a significant difference in global mean DNAme level by sex. Turning to individual CpG sites, we identified 162 DMPs across all autosomes that showed robust DNAme differences by placental sex with no evidence for cross-reactivity to the sex chromosomes. Functionally, these DMPs were enriched to be in or near genes associated with biological process gene ontology terms related to chemokines or chemotaxis and immune function or epithelial barrier function.

Of the 162 sex-associated DMPs identified, >90% were more highly methylated in male placental samples than in female samples, and this trend held true at all biological and statistical thresholds considered. Interestingly, EWAS of somatic tissues have revealed the opposite pattern, across studies the majority of somatic sex-associated DMPs are more highly methylated in female samples (52). This has been reported in studies of blood (53, 54), buccal swab (53), prefrontal cortex (55), pancreatic islets (56), and also in a meta-analysis of 36 somatic tissues (52). It is interesting to see the opposite trend in placenta (more DMPs with higher male DNAme), but perhaps not surprising, as this pattern was previously reported in a study of placental autosomal DNAme (13). Additionally, a study of placental DNAme by whole-genome oxidative bisulfite sequencing identified that male placentae are on the order of 1-2% more highly methylated overall than females (57); although we saw no significant difference in array-wide average DNAme by sex, this could be related to the uneven probe distribution of the 450K array, which are concentrated in functionally relevant areas (58). While the underlying cause of such a pattern in unclear, our investigation into a limited sample of placentae with a 45,X karyotype may suggest a role for X chromosome dosage. Studies of sex chromosome aneuploidies have revealed extensive influences of X chromosome dosage on DNAme profiles autosomal loci, for example in females affected by Turner syndrome and males affected by Klinefelter syndrome (59, 60). Additionally, as it has been proposed that X chromosome inactivation may be distinct (less complete) in the human placenta as compared to somatic tissues (61), It is possible that the placental inactive X may interact differently with autosomal loci than in somatic tissues.

In interpreting the biological significance of the 162 sex-associated DMPs, the genes overlapping these loci were enriched for biological process gene ontology terms related to chemokines and chemotaxis, as well as to the process of keratinization. This may suggest that the placenta mediates sex differential immune function and/or placental trophoblast structure or function during gestation, as genes from the KRT or keratin gene family are often used as cell-surface markers of placental trophoblasts (62), the most abundant placental cell type (63). Several genes from the ZNF family also overlapped DMPs and DMRs. *ZNF423* and *ZNF300*, specifically, overlap DMPs that are more highly methylated in male samples, and are both DNA-binding Krüppel-like C2H2 zinc finger transcription factors (64). *ZNF300* has been reported to be more highly expressed in female placentae in a study of first trimester conceptuses (16), this is consistent with the higher male DNAme in the *ZNF300* promoter we observe here (Figure 2). *ZNF423* was recently reported to regulate networks of gene co-expression (co-expression modules) in the human placenta that are conserved across gestation (15). Along with the *ENF1* gene, *ZNF423* regulated the most highly conserved placental co-expression module between humans and mice, suggesting the importance of *ZNF423* in the regulation of patterns of placental gene expression. To our knowledge, sex differences in placental DNAme of ZNF423 have not previously been reported, nor were sex differences in the *ZNF423* co-expression module reported. The sex-specific DNAme observed in this study across *ZNF423* could suggest that the conserved placental co-expression module identified by Buckberry et al. may be regulated in a sex-specific manner. For the plots shown in Figure 2, the location of all CpG sites shown aligned with the RefGene and ChromHMM tracks from the UCSC Human Genome Browser (65) are available in Supplementary Figure 3.

To understand the extent to which our DMPs were related to sex differences in placental gene expression, we investigated placental microarray expression data for the 65 genes overlapping the 162 DMPs identified. Although 12% of the 65 genes overlapping the 162 DMPs showed sex-specific placental expression, the majority were not significantly differentially expressed by placental sex. This is may be related to the small sample size of the gene expression cohort utilized (n=34), the role of additional factors beyond DNAme in regulating gene expression, and the possibility of alternative splicing and sex-specific isoform expression, which would not be captured in microarray analysis (66). Additionally, sex differences in DNAme at these 162 DMPs may be involved in regulating the expression of genes beyond those they overlap, which would not have been captured in this candidate gene expression analysis.

Both sex chromosome complement and relative sex hormone concentration can influence sex differences during gestation, as the conceptus harbors a sex chromosome complement and fetal testosterone begins to be produced by both sexes in the mid-first trimester (67). In the absence of amniotic fluid hormone measurements, we cannot comment extensively on the role fetal sex hormones play in establishing the DNAme profiles at these DMPs. However, DNAme profiles in female 45,X placental samples appeared more male-like in principal components analysis of the 162 DMPs, suggesting X chromosome complement may be associated with sex differences in placental autosomal DNAme. A further link between DMP DNAme profiles and the sex chromosomes was found in the enrichment for overlap with KAISO protein binding motifs. KAISO is a transcription factor encoded by the X-linked *ZBTB33* gene, and has been reported to repress gene expression by binding methylated DNA (68). The fact that *ZBTB33* is X-linked may imply the existence of interactions between the sex chromosomal and autosomal loci in the placenta. Furthermore, we found no association of differential DNAme with nearby ER or AR binding sites, making it less likely that hormone effects underly these differences.

Another outcome of our principal components analysis was the ability to observe associations between DNAme profiles at the 162 DMPs and sample demographic characteristics. Interestingly, we found that across the first principal component, in male samples increased maternal age was significantly associated with falling toward the male extreme of the continuum, while amongst females increased birthweight standard deviation was associated with the female end of the continuum. While maternal age has been positively associated with increased risk of preeclampsia development, we are not aware of sex differences in preeclampsia risk by maternal age (69). Conversely, birthweight standard deviation is a metric that is calculated using sex- and gestational age-adjusted growth curves, and as such is independent of both sex and gestational age. Although birthweight standard deviation was not expected to and did not differ significantly by sex in these cases, within the female samples a higher birthweight standard deviation was associated with those samples localizing toward the female extreme of PC1. While there are known sex differences in average birthweight, with males tending to be born heavier than females, to our knowledge this is the first report suggesting that placental molecular features may interact with intra-sex birthweight distributions.

In comparing the DMPs discovered in this study to findings previously reported in the human placenta (13, 14) we observed limited overlap, although all of the 85 DMPs from our study overlapped with previous reports were differentially methylated in the same direction by sex as previously reported. Limited overlap may partially relate to cohort size, as the cohort used in this study is larger than any used previously (341 samples versus 62 and 84 samples), increasing our power to detect true positive sex differences. Despite imperfect overlap with previous studies, we observed a high degree of DMP reproducibility between our discovery and replication cohorts, suggesting that the 162 DMPs identified here show consistent sex differences in placental autosomal DNA.

We acknowledge several limitations of our findings. First, because the discovery cohort utilized is largely of European and East Asian ancestry, and the replication dataset is comprised exclusively of European ancestry samples (21), our results may not generalize to other ancestral populations. This is a limitation applying to nearly every large-scale epigenome or genome-wide association study (70, 71), and inclusion of samples of diverse ancestry should be considered in the construction of future cohorts. Second, although enriched for coverage of functional genomic regions and RefSeq genes, the Illumina 450K array does not provide coverage of all genomic CpGs, specifically in non-coding regions. Higher-resolution technologies such as the Illumina EPIC array or whole-genome bisulfite sequencing can address this limitation. Further, we could not directly examine the relationship between placental DNAme and fetal sex hormone levels in amniotic fluid. We acknowledge that by term, both sex chromosome complement and sex hormone levels have had ample opportunity to exert their effects, and thus we cannot disentangle which patterns of sex-specific DNAme observed may be related to each.

### PERSPECTIVES & SIGNIFICANCE

In summary, we find that autosomal sex differences in DNAme exist in the human placenta, and in contrast to somatic tissues the majority of placental autosomal sex-differentially methylated CpG sites are more highly methylated in male samples. Additionally, patterns of DNAme at these CpG sites suggest that male (XY) and female (XX) placenta vary continuously, rather than discretely, at these autosomal loci. These results are intended to establish a baseline for sex differences existing in the uncomplicated term placenta’s autosomal methylome, and we anticipate that they will be useful to contextualize results of analyses from the placentae of complicated pregnancies, especially those complications with sex-biased phenotypes such as preterm birth and early-onset preeclampsia

## CONCLUSIONS

Overall, our study reports high-confidence and large-effect size autosomal sex-associated DMPs in the human placenta, and identifies several key features of these loci that suggest their potential relevance to sex differences observed in normative and complicated human gestations. It remains to be determined how these patterns of sex-specific placental DNAme arise, and what their functional implications are. We hope our findings facilitate future investigation of sex differences in placental molecular features as a means to investigating the causes and consequences of sex differences in pregnancy.

## DECLARATIONS

### Ethics approval and consent to participate

Ethics approval for use of human research subjects in this study was obtained from the University of British Columbia/Children’s and Women’s Health Centre of British Columbia Research Ethics Board (H18-01695). Informed written consent was obtained from all study participants.

## Consent for publication

Not applicable.

## Availability of data and materials

All datasets used are publicly available via the Gene Expression Omnibus at the indicated accession numbers (https://www.ncbi.nlm.nih.gov/geo/).

## Authors’ contributions

AMI contributed to study design, data preparation, and performed data analysis, interpretation, and drafted the manuscript. VY contributed to study design and prepared the datasets that compose the discovery cohort. CK contributed to study design and analysis. WPR, CJB, and AMM conceived of and supported the study, and contributed to data analysis and interpretation of the results. All authors read and provided critical feedback on the manuscript, and approved the final version.

## Acknowledgements

We thank all cohort owners and the scientific community for their commitment to making scientific data publicly available, and we thank all study participants for the generous donation of samples. We acknowledge members of the Robinson lab for thoughtful discussion and feedback on the analysis and manuscript, especially Giulia F. Del Gobbo, Dr. Maria Peñaherrera A., and Dr. Johanna Schuetz.

## ADDITIONAL FILES

**Supplementary Figures (suppfigures.pdf)**

Title: Supplementary figure files

Description: Supplementary figures 1-4 with corresponding titles and figure captions.

---

**Supplementary Table 1 (supptable_1.xlsx)**

Title: Results of linear modelling for all 324,104 autosomal CpGs tested

Description: Linear modelling statistics for sex differential methylation analysis at all 324,104 autosomal CpG sites in the filtered dataset.

**Supplementary Table 2 (supptable_2.xlsx)**

Title: Table of significant placental autosomal sex-associated DMRs

Description: Summary statistics and genomic locations of all signficant sex-associated DMRs

identified.

**Supplementary Table 3 (supptable_3.xlsx)**

Title: Table of significant placental autosomal sex-associated DVPs

Description: Summary statistics and genomic locations of differential variability analysis for all significant sex-associated DVPs.

## ABBREVIATIONS

450K: Illumina HumanMethylation450 Array

ANOVA: analysis of variance

AR: androgen receptor

BLAST: basic local alignment search tool

BMIQ: Beta-Mixture Quantile Normalization

CpG: cytosine-guanine dinucleotide

DMP: differentially methylated position (1 CpG)

DMR: differentially methylated genomic region (>1 CpG)

DNAme: DNA methylation

DVP: differentially variably methylated position (1 CpG)

ER: estrogen receptor

FDR: Benjamini-Hochberg false discovery rate

LGA: large birthweight for gestational age

PC: principal component

PCA: principal components analysis

PlaNET: Placental DNAme Elastic Net Ethnicity Tool

SD: standard deviation

SGA: small birthweight for gestational age

XCI: X chromosome inactivation

## REFERENCES

1. Arnold AP. A general theory of sexual differentiation. J Neurosci Res. 2017 02;95(1–2):291–300.

2. Sandman CA, Glynn LM, Davis EP. Is there a viability–vulnerability tradeoff? Sex differences in fetal programming. Journal of Psychosomatic Research. 2013 Oct 1;75(4):327–35.

3. Challis J, Newnham J, Petraglia F, Yeganegi M, Bocking A. Fetal sex and preterm birth. Placenta. 2013 Feb 1;34(2):95–9.

4. Di Renzo GC, Rosati A, Sarti RD, Cruciani L, Cutuli AM. Does fetal sex affect pregnancy outcome? Gender Medicine. 2007 Mar 1;4(1):19–30.

5. Clifton VL. Review: Sex and the Human Placenta: Mediating Differential Strategies of Fetal Growth and Survival. Placenta. 2010 Mar 1;31:S33–9.

6. Broere-Brown ZA, Adank MC, Benschop L, Tielemans M, Muka T, Gonçalves R, et al. Fetal sex and maternal pregnancy outcomes: a systematic review and meta-analysis. Biol Sex Differ [Internet]. 2020 May 11;11. Available from: https://www.ncbi.nlm.nih.gov/pmc/articles/PMC7216628/

7. Bale TL. The placenta and neurodevelopment: sex differences in prenatal vulnerability. Dialogues Clin Neurosci. 2016 Dec;18(4):459–64.

8. Rosenfeld CS. Sex-Specific Placental Responses in Fetal Development. Endocrinology. 2015 Oct 1;156(10):3422–34.

9. Sharp AJ, Stathaki E, Migliavacca E, Brahmachary M, Montgomery SB, Dupre Y, et al. DNA methylation profiles of human active and inactive X chromosomes. Genome Res [Internet]. 2011 Aug 23 [cited 2018 Apr 23]; Available from: http://genome.cshlp.org/content/early/2011/08/23/gr.112680.110

10. Cotton AM, Price EM, Jones MJ, Balaton BP, Kobor MS, Brown CJ. Landscape of DNA methylation on the X chromosome reflects CpG density, functional chromatin state and X-chromosome inactivation. Hum Mol Genet. 2015 Mar 15;24(6):1528–39.

11. Blair JD, Price EM. Illuminating Potential Technical Artifacts of DNA-Methylation Array Probes. Am J Hum Genet. 2012 Oct 5;91(4):760–2.

12. Chen Y, Choufani S, Grafodatskaya D, Butcher DT, Ferreira JC, Weksberg R. Cross-Reactive DNA Microarray Probes Lead to False Discovery of Autosomal Sex-Associated DNA Methylation. Am J Hum Genet. 2012 Oct 5;91(4):762–4.

13. Martin E, Smeester L, Bommarito PA, Grace MR, Boggess K, Kuban K, et al. Sexual epigenetic dimorphism in the human placenta: implications for susceptibility during the prenatal period. Epigenomics. 2017 Mar;9(3):267–78.

14. Mayne BT, Leemaqz SY, Smith AK, Breen J, Roberts CT, Bianco-Miotto T. Accelerated placental aging in early onset preeclampsia pregnancies identified by DNA methylation. Epigenomics. 2016 Nov 29;9(3):279–89.

15. Buckberry S, Bianco-Miotto T, Bent SJ, Dekker GA, Roberts CT. Integrative transcriptome meta-analysis reveals widespread sex-biased gene expression at the human fetal–maternal interface. Mol Hum Reprod. 2014 Aug;20(8):810–9.

16. Gonzalez TL, Sun T, Koeppel AF, Lee B, Wang ET, Farber CR, et al. Sex differences in the late first trimester human placenta transcriptome. Biology of Sex Differences. 2018 Jan 15;9:4.

17. Wilson SL, Leavey K, Cox B, Robinson WP. The value of DNA methylation profiling in characterizing preeclampsia and intrauterine growth restriction. bioRxiv. 2017 Jun 18;151290.

18. Konwar C, Price EM, Wang LQ, Wilson SL, Terry J, Robinson WP. DNA methylation profiling of acute chorioamnionitis-associated placentas and fetal membranes: insights into epigenetic variation in spontaneous preterm births. Epigenetics & Chromatin. 2018 Oct 29;11(1):63.

19. Roifman M, Choufani S, Turinsky AL, Drewlo S, Keating S, Brudno M, et al. Genome-wide placental DNA methylation analysis of severely growth-discordant monochorionic twins reveals novel epigenetic targets for intrauterine growth restriction. Clin Epigenetics [Internet]. 2016 Jun 21 [cited 2020 Apr 15];8. Available from: https://www.ncbi.nlm.nih.gov/pmc/articles/PMC4915063/

20. Maccani JZ, Koestler DC, Houseman EA, Marsit CJ, Kelsey KT. Placental DNA methylation alterations associated with maternal tobacco smoking at the RUNX3 gene are also associated with gestational age. Epigenomics. 2013 Dec;5(6):619–30.

21. Green BB, Karagas MR, Punshon T, Jackson BP, Robbins DJ, Houseman EA, et al. Epigenome-Wide Assessment of DNA Methylation in the Placenta and Arsenic Exposure in the New Hampshire Birth Cohort Study (USA). Environ Health Perspect. 2016;124(8):1253– 60.

22. Everson TM, Punshon T, Jackson BP, Hao K, Lambertini L, Chen J, et al. Cadmium-Associated Differential Methylation throughout the Placental Genome: Epigenome-Wide Association Study of Two U.S. Birth Cohorts. Environ Health Perspect. 2018 22;126(1):017010.

23. Martin E, Ray PD, Smeester L, Grace MR, Boggess K, Fry RC. Epigenetics and Preeclampsia: Defining Functional Epimutations in the Preeclamptic Placenta Related to the TGF-β Pathway. PLOS ONE. 2015 Oct 28;10(10):e0141294.

24. Paquette AG, Houseman EA, Green BB, Lesseur C, Armstrong DA, Lester B, et al. Regions of variable DNA methylation in human placenta associated with newborn neurobehavior. Epigenetics. 2016 Jul 1;11(8):603–13.

25. Leavey K, Wilson SL, Bainbridge SA, Robinson WP, Cox BJ. Epigenetic regulation of placental gene expression in transcriptional subtypes of preeclampsia. Clin Epigenetics [Internet]. 2018 Mar 2 [cited 2018 Jun 12];10. Available from: https://www.ncbi.nlm.nih.gov/pmc/articles/PMC5833042/

26. Hanna CW, Peñaherrera MS, Saadeh H, Andrews S, McFadden DE, Kelsey G, et al. Pervasive polymorphic imprinted methylation in the human placenta. Genome Res. 2016 Jun;26(6):756–67.

27. Wilson SL, Leavey K, Cox BJ, Robinson WP. Mining DNA methylation alterations towards a classification of placental pathologies. Hum Mol Genet. 2018 Jan 1;27(1):135–46.

28. Yuan V, Price EM, Del Gobbo G, Mostafavi S, Cox B, Binder AM, et al. Accurate ethnicity prediction from placental DNA methylation data. Epigenetics & Chromatin. 2019 Aug 9;12(1):51.

29. Price EM, Robinson WP. Adjusting for Batch Effects in DNA Methylation Microarray Data, a Lesson Learned. Front Genet [Internet]. 2018 [cited 2018 Apr 17];9. Available from: https://www.frontiersin.org/articles/10.3389/fgene.2018.00083/full

30. Heiss JA, Just AC. Identifying mislabeled and contaminated DNA methylation microarray data: an extended quality control toolset with examples from GEO. Clinical Epigenetics. 2018 Jun 1;10(1):73.

31. Chen Y, Lemire M, Choufani S, Butcher DT, Grafodatskaya D, Zanke BW, et al. Discovery of cross-reactive probes and polymorphic CpGs in the Illumina Infinium HumanMethylation450 microarray. Epigenetics. 2013 Feb 1;8(2):203–9.

32. Price ME, Cotton AM, Lam LL, Farré P, Emberly E, Brown CJ, et al. Additional annotation enhances potential for biologically-relevant analysis of the Illumina Infinium HumanMethylation450 BeadChip array. Epigenetics & Chromatin. 2013;6(1):4.

33. Edgar RD, Jones MJ, Robinson WP, Kobor MS. An empirically driven data reduction method on the human 450K methylation array to remove tissue specific non-variable CpGs. Clinical Epigenetics. 2017 Feb 2;9(1):11.

34. Aryee MJ, Jaffe AE, Corrada-Bravo H, Ladd-Acosta C, Feinberg AP, Hansen KD, et al. Minfi: a flexible and comprehensive Bioconductor package for the analysis of Infinium DNA methylation microarrays. Bioinformatics. 2014 May 15;30(10):1363–9.

35. Teschendorff AE, Marabita F, Lechner M, Bartlett T, Tegner J, Gomez-Cabrero D, et al. A beta-mixture quantile normalization method for correcting probe design bias in Illumina Infinium 450 k DNA methylation data. Bioinformatics. 2013 Jan 15;29(2):189–96.

36. Smit A, Hubley R. RepeatMasker Open-4.0 [Internet]. 2013. Available from: http://www.repeatmasker.org

37. Ritchie ME, Phipson B, Wu D, Hu Y, Law CW, Shi W, et al. limma powers differential expression analyses for RNA-sequencing and microarray studies. Nucleic Acids Res. 2015 Apr 20;43(7):e47–e47.

38. Phipson B, Maksimovic J, Oshlack A. missMethyl: an R package for analyzing data from Illumina’s HumanMethylation450 platform. Bioinformatics. 2016 Jan 15;32(2):286–8.

39. Kulakovskiy IV, Vorontsov IE, Yevshin IS, Sharipov RN, Fedorova AD, Rumynskiy EI, et al. HOCOMOCO: towards a complete collection of transcription factor binding models for human and mouse via large-scale ChIP-Seq analysis. Nucleic Acids Res. 2018 Jan 4;46(Database issue):D252–9.

40. Bailey TL, Machanick P. Inferring direct DNA binding from ChIP-seq. Nucleic Acids Res. 2012 Sep 1;40(17):e128.

41. Bailey TL, Boden M, Buske FA, Frith M, Grant CE, Clementi L, et al. MEME Suite: tools for motif discovery and searching. Nucleic Acids Res. 2009 Jul 1;37(suppl_2):W202–8.

42. Wilson S, Qi J, Filipp FV. Refinement of the androgen response element based on ChIP-Seq in androgen-insensitive and androgen-responsive prostate cancer cell lines. Scientific Reports. 2016 Sep 14;6(1):32611.

43. Grober OM, Mutarelli M, Giurato G, Ravo M, Cicatiello L, De Filippo MR, et al. Global analysis of estrogen receptor beta binding to breast cancer cell genome reveals an extensive interplay with estrogen receptor alpha for target gene regulation. BMC Genomics. 2011 Jan 14;12(1):36.

44. Leavey Katherine, Benton Samantha J., Grynspan David, Kingdom John C., Bainbridge Shannon A., Cox Brian J. Unsupervised Placental Gene Expression Profiling Identifies Clinically Relevant Subclasses of Human Preeclampsia. Hypertension. 2016 Jul 1;68(1):137– 47.

45. Peters TJ, Buckley MJ, Statham AL, Pidsley R, Samaras K, Lord RV, et al. De novo identification of differentially methylated regions in the human genome. 2015;16.

46. Feinberg AP, Irizarry RA. Stochastic epigenetic variation as a driving force of development, evolutionary adaptation, and disease. PNAS. 2010 Jan 26;107(suppl 1):1757–64.

47. Price EM, Cotton AM, Peñaherrera MS, McFadden DE, Kobor MS, Robinson W. Different measures of “genome-wide” DNA methylation exhibit unique properties in placental and somatic tissues. Epigenetics. 2012 Jun 1;7(6):652–63.

48. Konwar C, Del Gobbo G, Yuan V, Robinson WP. Considerations when processing and interpreting genomics data of the placenta. Placenta. 2019 Sep;84:57–62.

49. Costanzo V, Bardelli A, Siena S, Abrignani S. Exploring the links between cancer and placenta development. Open Biol [Internet]. 2018 Jun 27 [cited 2021 Jan 25];8(6). Available from: https://www.ncbi.nlm.nih.gov/pmc/articles/PMC6030113/

50. Kane MD, Jatkoe TA, Stumpf CR, Lu J, Thomas JD, Madore SJ. Assessment of the sensitivity and specificity of oligonucleotide (50mer) microarrays. Nucleic Acids Res. 2000 Nov 15;28(22):4552–7.

51. Blair JD, Yuen RKC, Lim BK, McFadden DE, von Dadelszen P, Robinson WP. Widespread DNA hypomethylation at gene enhancer regions in placentas associated with early-onset pre-eclampsia. Mol Hum Reprod. 2013 Oct;19(10):697–708.

52. McCarthy NS, Melton PE, Cadby G, Yazar S, Franchina M, Moses EK, et al. Meta-analysis of human methylation data for evidence of sex-specific autosomal patterns. BMC Genomics [Internet]. 2014 Nov 18 [cited 2018 Apr 13];15(1). Available from: https://www.ncbi.nlm.nih.gov/pmc/articles/PMC4255932/

53. Singmann P, Shem-Tov D, Wahl S, Grallert H, Fiorito G, Shin S-Y, et al. Characterization of whole-genome autosomal differences of DNA methylation between men and women. Epigenetics Chromatin [Internet]. 2015 Oct 19 [cited 2018 Apr 18];8. Available from: https://www.ncbi.nlm.nih.gov/pmc/articles/PMC4615866/

54. Yousefi P, Huen K, Davé V, Barcellos L, Eskenazi B, Holland N. Sex differences in DNA methylation assessed by 450 K BeadChip in newborns. BMC Genomics [Internet]. 2015 Nov 9 [cited 2018 Apr 18];16. Available from: https://www.ncbi.nlm.nih.gov/pmc/articles/PMC4640166/

55. Xu H, Wang F, Liu Y, Yu Y, Gelernter J, Zhang H. Sex-biased methylome and transcriptome in human prefrontal cortex. Hum Mol Genet. 2014 Mar 1;23(5):1260–70.

56. Hall E, Volkov P, Dayeh T, Esguerra JLS, Salö S, Eliasson L, et al. Sex differences in the genome-wide DNA methylation pattern and impact on gene expression, microRNA levels and insulin secretion in human pancreatic islets. Genome Biol [Internet]. 2014 [cited 2018 Apr 24];15(12). Available from: https://www.ncbi.nlm.nih.gov/pmc/articles/PMC4256841/

57. Gong S, Johnson MD, Dopierala J, Gaccioli F, Sovio U, Constância M, et al. Genome-wide oxidative bisulfite sequencing identifies sex-specific methylation differences in the human placenta. Epigenetics. 2018 Jan 29;0(0):1–12.

58. Sandoval J, Heyn H, Moran S, Serra-Musach J, Pujana MA, Bibikova M, et al. Validation of a DNA methylation microarray for 450,000 CpG sites in the human genome. Epigenetics. 2011 Jun;6(6):692–702.

59. Trolle C, Nielsen MM, Skakkebæk A, Lamy P, Vang S, Hedegaard J, et al. Widespread DNA hypomethylation and differential gene expression in Turner syndrome. Scientific Reports. 2016 Sep 30;6:34220.

60. Skakkebæk A, Nielsen MM, Trolle C, Vang S, Hornshøj H, Hedegaard J, et al. DNA hypermethylation and differential gene expression associated with Klinefelter syndrome. Scientific Reports. 2018 Sep 13;8(1):13740.

61. Cotton AM, Avila L, Penaherrera MS, Affleck JG, Robinson WP, Brown CJ. Inactive X chromosome-specific reduction in placental DNA methylation. Hum Mol Genet. 2009 Oct 1;18(19):3544–52.

62. Gauster M, Blaschitz A, Siwetz M, Huppertz B. Keratins in the human trophoblast. Histol Histopathol. 2013 Jul;28(7):817–25.

63. Yuan V, Hui D, Yin Y, Peñaherrera MS, Beristain AG, Robinson WP. Cell-specific characterization of the placental methylome. BMC Genomics. 2021 Jan 6;22(1):6.

64. Cassandri M, Smirnov A, Novelli F, Pitolli C, Agostini M, Malewicz M, et al. Zinc-finger proteins in health and disease. Cell Death Discov. 2017;3:17071.

65. Kent WJ, Sugnet CW, Furey TS, Roskin KM, Pringle TH, Zahler AM, et al. The human genome browser at UCSC. Genome Res. 2002 Jun;12(6):996–1006.

66. Karlebach G, Veiga DFT, Mays AD, Kesarwani AK, Danis D, Kararigas G, et al. The impact of sex on alternative splicing.: 24.

67. Künzig HJ, Meyer U, Schmitz-Roeckerath B, Broer KH. Influence of fetal sex on the concentration of amniotic fluid testosterone: Antenatal sex determination? Arch Gynak. 1977;223(2):75–84.

68. Yoon H-G, Chan DW, Reynolds AB, Qin J, Wong J. N-CoR Mediates DNA Methylation-Dependent Repression through a Methyl CpG Binding Protein Kaiso. Molecular Cell. 2003 Sep 1;12(3):723–34.

69. Cavazos-Rehg PA, Krauss MJ, Spitznagel EL, Bommarito K, Madden T, Olsen MA, et al. Maternal age and risk of labor and delivery complications. Matern Child Health J. 2015 Jun;19(6):1202–11.

70. Popejoy AB, Fullerton SM. Genomics is failing on diversity. Nature. 2016 Oct 12;538(7624):161–4.

71. Kessler MD, Yerges-Armstrong L, Taub MA, Shetty AC, Maloney K, Jeng LJB, et al. Challenges and disparities in the application of personalized genomic medicine to populations with African ancestry. Nature Communications. 2016 Oct 11;7(1):12521.

