## Supplementary Figures for "A cross-cohort analysis of autosomal DNA methylation sex differences in the term placenta"

**Supplementary Figure 1.** *PlaNET-estimated ancestry of all replication cohort samples (GSE71678).* Scatterplot PlaNET ancestry coordinates 2 and 3, reflecting East Asian and European ancestry, for all samples in the replication cohort GSE71678 (n=293). Estimated ancestry in this cohort is extremely homogeneous (predominantly European) as indicated by all 293 samples falling in the top left corner of the plot. Samples are coloured by PlaNET-inferred ancestry, Blue represents likely European ancestry, Red represents ambiguous ancestry (admixed).

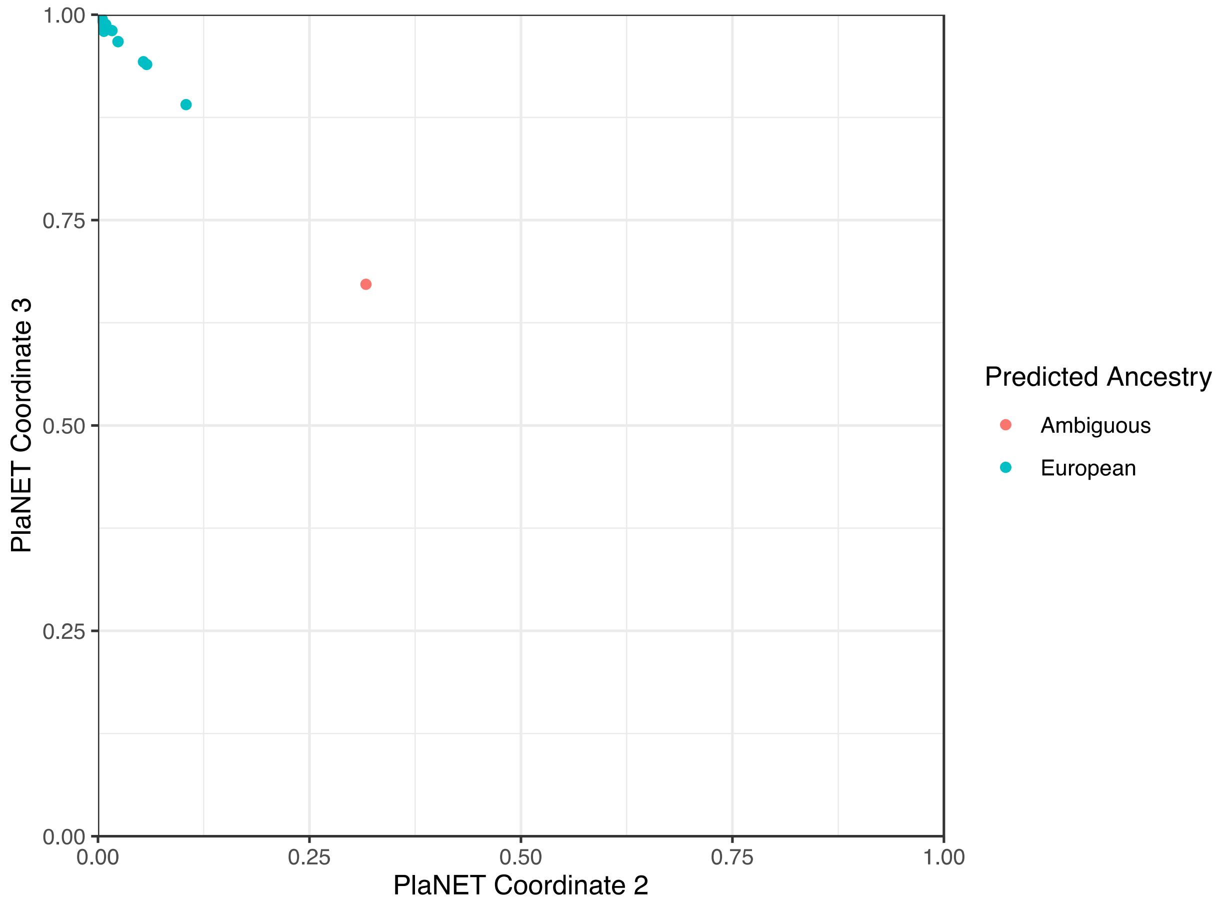

**Supplementary Figure 2.** Potential impact on sex-associated DNAme of putative X chromosomal cross-hybridization  of probe cg02325951, for which the intended genomic target is chr14: 89878619-89878668.

β

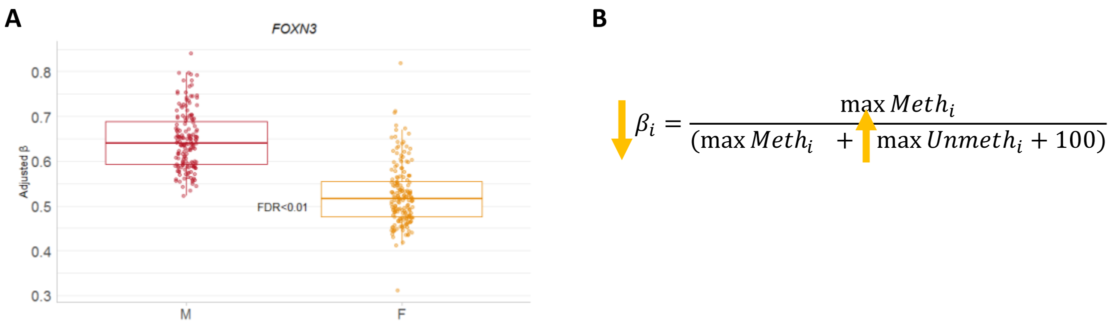

**Supplementary Figure 3**. UCSC Genome Browser plots of regions shown in Figure 2 for (A) ZNF300, (B) ZNF423, and (C) SPON1, including the location of all CpGs plotted with reference to the UCSC gene tracks and ChromHMM states. This figure was created using resources provided by http://genome.ucsc.edu.

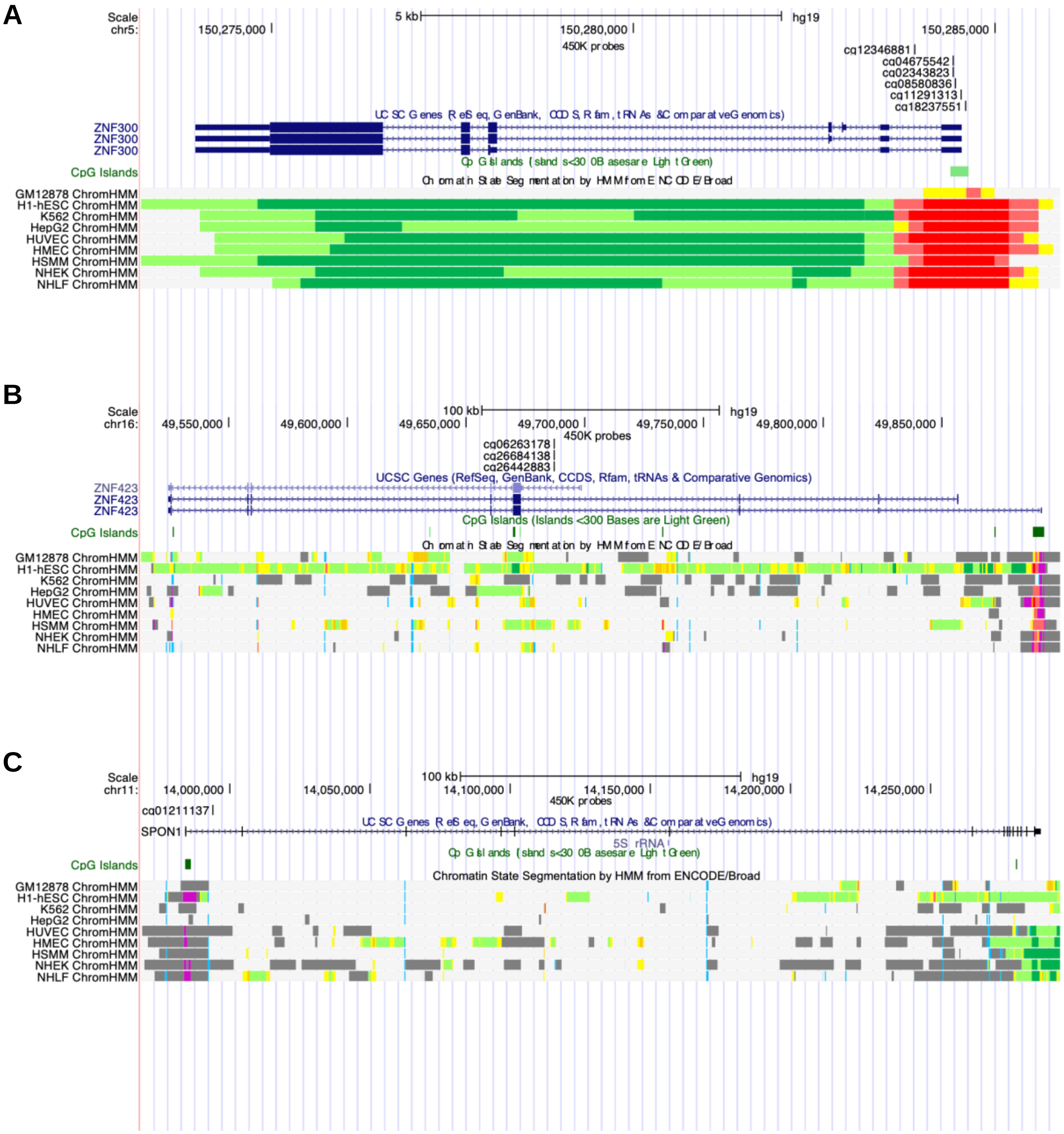

**Supplementary Figure 4**. Principal components analysis on 162 autosomal DMPs (A) Scatterplot of PC1 versus PC2 for the 34-sample subset of the discovery cohort for which extended demographic information was available. (B) Projection of three 45,X placental samples in the principal components space associated with DNAme patterns at the top 162 DMPs in all samples from the discovery cohort.

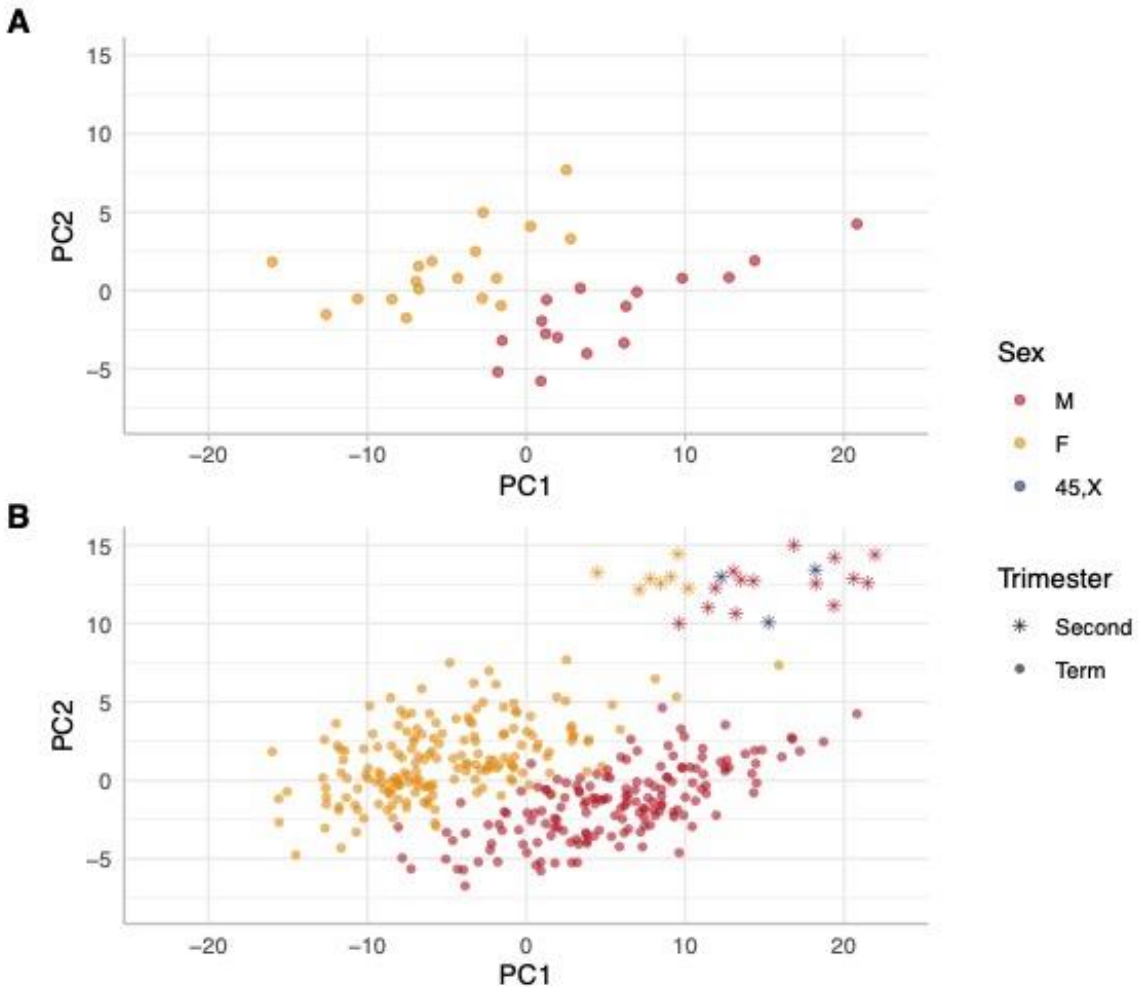
